# Clean-PIE: a novel strategy for efficiently constructing precise circRNA with thoroughly minimized immunogenicity to direct potent and durable protein expression

**DOI:** 10.1101/2022.06.20.496777

**Authors:** Zonghao Qiu, Qiangbo Hou, Yang Zhao, Jiafeng Zhu, Mengting Zhai, Daolei Li, Yi Li, Chunxi Liu, Na Li, Yifei Cao, Jiali Yang, Zhenhua Sun, Chijian Zuo

## Abstract

Translatable circular RNAs (circRNAs) are emerging as a crucial molecular format for transient protein expression, with high potential to be an alternative for linear mRNA to reshape the landscape of mRNA pharmaceutical industry. Canonical *Anabaena* permuted intron-exon (*Ana* PIE) format that developed by ORNA is an efficient method for RNA circularization, and the engineered circRNAs direct supreme protein expression in eukaryotic cells. However, recent studies revealed that this method may unavoidably result in a remain of immunogenicity in the circRNA products, albeit after thorough RNA purification. In the current study, we develop a novel strategy for efficient generation of circRNA, via the permuted T4Td introns mediated autocatalytically ribozymatic reaction mediated ligation of the flanking segment sequences that concealing in ORF or translation initiation sequence (normally equal to IRES). This strategy universally realizes around 90% circularization effectivity, and the circRNA products can be purified to around 90% purity by our new purification method, and presented thoroughly minimized immunogenicity, thus is termed “Clean-PIE”. The purified circRNAs are found to direct potent and durable expression of various proteins in vitro and in vivo. The partly purified Fluc circRNA by HPLC-SEC was found to direct Fluc expression in muscle for no less than 20 days. The highly purified circRNA exhibits much stronger protein expression in vitro and in vivo, and presumed a longer duration. Additionally, the scale-up of RNA circularization with the RNA precursors from 1 L transcription revealed high circularization effectivity (around 90%) and a high productivity of the final circRNA products. Collectively, Clean-PIE is a novel circRNA platform that possesses high circularization effectivity, enabled high RNA purity and thoroughly minimized immunogenicity, as well as scaling-up accessibility and directing extreme durability of protein expression, thus has the potential to develop advanced RNA vaccines and therapeutics in pharmaceutical industrial scale.

## Introduction

Circular RNAs (circRNAs), as single-stranded and covalently closed RNAs, were recently identified as a universal class of non-coding RNA in eukaryotic cells^1,2^. Researchers have demonstrated that circular RNAs with an internal ribosome entry site (IRES) can be translated both *in vitro* and *in vivo* ^3–5^. Furthermore, a scalable method for producing continuously translating circular mRNA has been proposed^6,7^. According to Wesselhoeft et al., circulating RNAs, produced massively as a result of *in vitro* transcription reactions, can serve as stable and potent alternatives to linear mRNA for translation^8^. Contrary to the linear form of mRNA, the circular form of mRNA has unique properties and additional capabilities. For instance, circular RNA exhibits higher stability and longer half-life than linear mRNA, thus may result in stronger and more durable protein expression in eucaryotic cells. Furthermore, the in vitro manufacturing of translatable circular mRNA is much simpler than linear mRNA, due to the replacement of sophisticated capping, polyA tail adding and modified nucleoside adding reactions^9^ by a single step circularization.

Albeit the immunogenicity of high-purity synthetic circRNAs was found diminished compared to unmodified linear mRNA^10,11^, further validations of the immunogenicity of circular RNAs are necessary. Recently, researchers found that the circRNA produced by the *Anabaena* permuted intron-exon (PIE) or the *Thymidylate* synthase (TD) PIE have significantly higher immunogenicity, compared with the one generated by T4 ligase^12^. The culprit was charged with extraneous fragments, exactly the long exon1-exon2 (E1-E2) junction that induced by Ana PIE and td PIE^12^. Thus, circRNAs generated by the circularization strategy that developed by Weesselhoeft et al. in Daniel. G. Anderson group or ORNA unavoidably possesses the remain of immunogenicity. This induction of extraneous fragments and the resultant immunogenicity can be avoided by using the T4 ligase method, however, the relative low circularization efficiency and the requirement for splints discourage the use of this method for high yield circRNA production. Moreover, transcripts usually exhibit highly heterogeneous 3’ ends, which renders the T4 ligase method for producing corrected-size circRNAs^13–15^. For these reasons, a more efficient, accurate, and immunogenicity thoroughly diminishing method for RNA circularization is required.

In the current study, our strategy presents a novel method for producing precise translatable circular RNA with a minimal-sized endogenous E1-E2 segment sequences concealing within a coding sequence, or within a translation initiation sequence, which is normally an internal ribosome entry site (IRES) element. In our RNA circularization strategy, permuted T4Td flanking introns are employed for autocatalytically ribozyme mediated splicing and circularization, and the flanking E1 and E2 sequences concealing in ORF or IRES are ligated and circRNAs generated, and free-style introns are generated as side products. This circularization strategy was proven to efficiently circularize various RNA precursors to generate circRNAs with the effectivity around 90%. CircRNAs generated by this method are highly translatable and mediated durable protein expressions in vitro and in vivo. Moreover, circRNAs that generated by this strategy possess thoroughly diminishing immunogenicity, compared with the one generated by ORNA strategy, probably due to thoroughly avoiding the involvement of extraneous fragments in circRNAs. Therefore, this circularization strategy is termed “Clean-PIE”. Importantly, the partly purified circRNA that encodes Fluc was found to express in muscle for at 20 days, suggesting an extreme stability and durability of protein expression possessed by this kind of RNA. The highly purified circRNA with around 90% purity by our new HPLC purification method improves the protein expression level of circRNA in vitro and in vivo, presumed to greatly extend the durability of in vivo expression. Furthermore, large-scale manufacturing of Fluc circRNAs up to 1 L of transcription was performed, around 90% circularization effectivity, and around 90% purity, as well as high productivity were realized. Collectively, Clean-PIE is a circRNA platform with universal high circularization effectivity, high-purity circRNA generation, thoroughly minimized immunogenicity, scale-up accessibility, and directing potent and extremely durable protein expressions, thus has the potential for the development of advanced RNA vaccines and therapeutics, and sufficient for manufacturing in pharmaceutical industry.

## Results

### To obtain precise circRNAs, conceal the T4td PIE splicing site in the ORF with codon optimization

Previously, several amounts of PIE-based RNA circularization strategies have been reported^8,11,16^. CircRNAs can be synthesized *in vitro* by circularization reaction using a transcription produced linear RNA percussor, in which two flanking transposed halves of a T4td or Anabaena derived split group I intron autocatalytically excise and the two flanking exons from T4td or Anabaena ligate in tandem transesterification reactions to form circRNAs^8,17,18^. It is important to note, however, that all of these PIE strategies introduce exogenous sequences into circularized RNAs, definitely from the retain of extraneous E1-E2 sequences. To synthesize precise, effective, and immunogenicity-free circRNAs as potential RNA vaccines and therapeutics, we attempted to reduce the introduced extraneous sequences in the *in vitro* synthetic circRNAs. Recently, Rausch JW *et al*. proved that the sequences of E1 and E2 for RNA ligation are not obligatory to the natural sequences of exons, however, can be a set of variants and high circularization effectivity maintained.

In this report, a design principle for variable E1 and E2 was discovered and confirmed in the model of constructing circPVT1, a 410 nt circular non-coding RNA^19^. Here, we attempted to extend this circRNA construction principle to large-size, translatable circRNAs. However, when we try to use this method to produce a large translatable circRNA (1625bp), the yield of the final circRNA is quite low (data not shown). The most critical sequences of the E1-E2 for the ligation reaction is the domain that forms P1 of 5’ intron and P9 of 3’ intron, through Watson-Crick base pairs or wobble pairs (especially G-U pair). For instance, the most effective E1-E2 sequences for T4td PIE is the TTGGGTCT, which forms P1 and part of P9 with internal short guide (IG) sequence (Figure. 1A and 1B) in *td* group I intron of bacteriophage T4 (T4td). We hypothesized to conceal these short E1 and E2 sequences into the ORF region of the circRNAs to avoid the introduction of exogenous T4td E1-E2 sequences, which may be the source of immunogenicity. As described by Rausch JW et al., except for the unchangeable ‘T’ ‘C’ site, which is critical for circularization reactions, the other sequences of E1-E2 are variable. Additionally, a script was developed for searching the optimal E1-E2 sequences, which is revealed by a scoring system (from 0 to 16 points of the score)^19^. Therefore, we speculate that the E1 and E2 sequences can be concealed in the coding sequences of ORF, implying that ORF can be segmented and flanked in the linear RNA precursor for ligation and circularization, as illustrated by the scheme in Figure.1A. Firstly, the given RNA sequences encoding eGFP was evaluated by the script provided by Rausch et al., indicating that the highest score is 9 points, with the segments as ‘TgGtGTCa’ and ‘aTGGaTCa’, respectively (Figure 1C). Next, the segment ‘aTGGaTCa’, as shown in Figure. 1C and 1D, located at the site ‘GATGGATCA’ coding for ‘Asp-Gly-Ser’, was selected as the segmenting site, and after nucleotides replacement, this segment was changed to ‘GATGGGTCT’ which is already quite closed to the E1E2 sequence of td PIE (‘TTGGGTCT’), without a change of the coding amino acids, due to the degeneracy of genetic codes. Next, the eGFP coding sequences was segmented as 3’ ORF and 5’ ORF, and flanked located besides the intron halves. The construction is shown in Figure. 1D. Circular RNA was produced as described in the method which was slightly modified by Wesselhoeft RA et al. As shown in Figure. 1E, the introns are spliced efficiently from the precursor, revealed by denaturing agarose gel analysis. To further confirm the functionality of this product, circular RNAs were transfected in 293T cells to monitor eGFP protein expression. Compared with *Ana* PIE constructed circRNA, circRNA that generated by our new circularization method resulted in unexpectedly higher eGFP protein expression.

**Figure 1:**
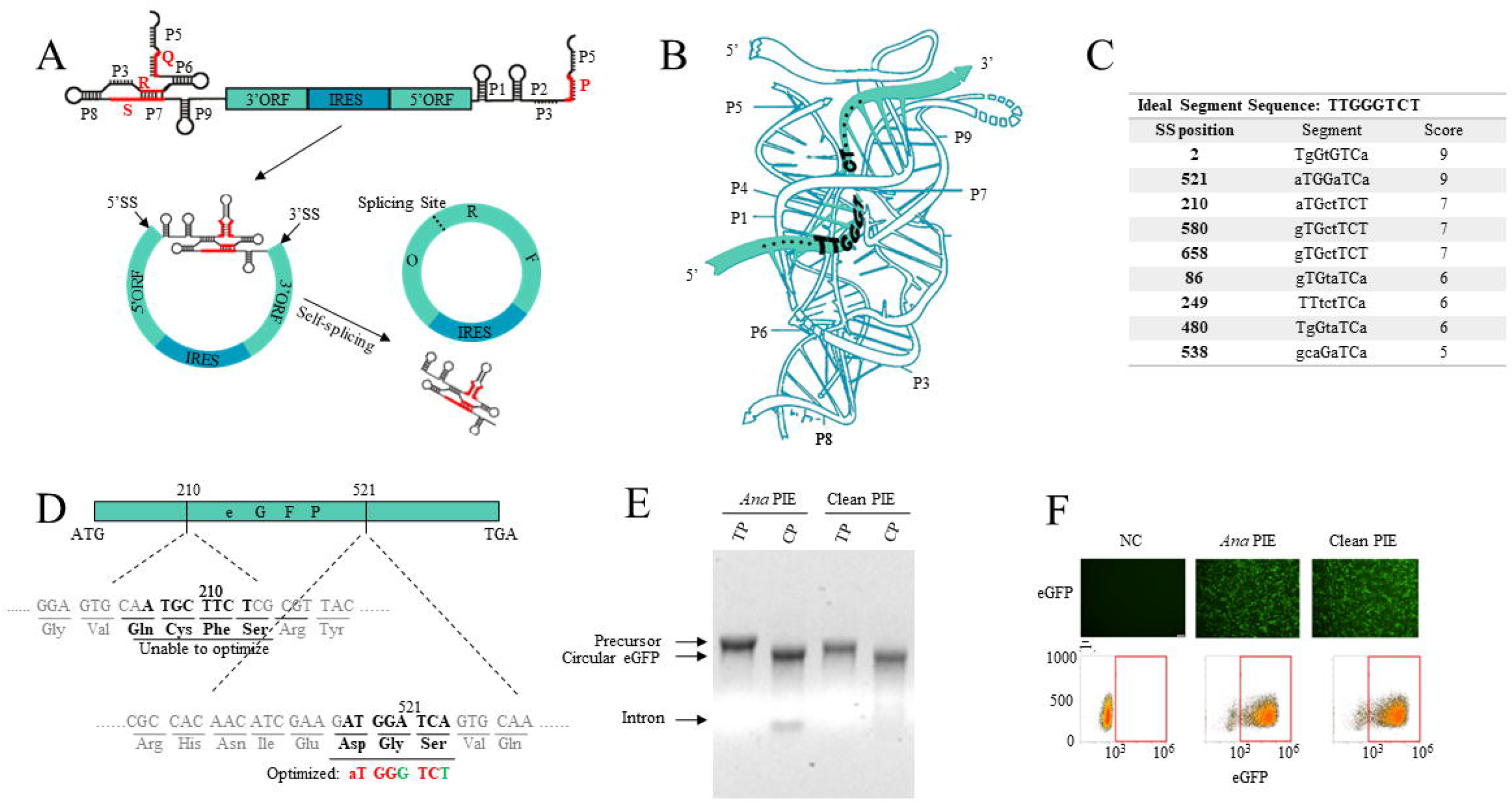
Concealing the splicing site of group I PIE in the ORF. A) A schematic diagram illustrating the design of permuted intron-exon constructs as well as the mechanisms of automatic self-splicing in group I introns. During truncation of the ORF, the splicing site is hidden within the ORF, and the split introns are flanked at the two sides. The conserved domains of group I intron are indicated as red color. B) Three-dimensional structure model of the td group I intron of the bacteriophage T4. The catalytic core of T4*td* is indicated by blue. The splicing site of T4td intron (TTGGGTCT) is indicated by light green. C) Sequences that are the closest match to T4td splicing site are listed in the table. D) Schematic showing how optimized splicing site is generated. Original sequences are presented in red, while optimized sequences are displayed in green. E) The agarose gel demonstrates that PIE circularization of the RNA is effective. TP: transcription products, CP: circularization products. F) Expression level detection of circular eGFP generated via different circularization strategies.

### Automatically finding the splicing site in ORF with a novel scoring system

The python script and scoring system that developed by Jason. W. Rausch et.al can be applied to estimate the efficiency of a E1-E2 segment for circularization, and an ideal segment can be identified and selected. In this study, we extended this principle to construct translatable circRNAs by concealing optimal E1-E2 segments in ORF for circularization. In the case of eGFP coding sequences in Figure.1, the segment sequence is already approximately identical to the original E1-E2 sequence of *td* PIE, after the manual replacement of two nucleotides. Next, we attempted to develop an algorithm assisted, automatically “nucleotide-replacement” system to optimize potential E1-E2 segments in ORF. Due to the degeneracy of genetic codes, RNA codons can be adapted to generate optimized E1-E2 sequences in ORF for effective circularization without a change of coding amino acids. Firstly, a sequence cutting system is designed. Take T4td PIE as an example, the minimal E1-E2 sequence ‘TTGGGTCT’ can be transformed into three different codon forms: ‘XTT-GGG-TCT’, ‘TTG-GGT-CTX’ and ‘XXT-TGG-GTC-TXX’, herein “X” represents a random nucleotide. Like this, the given amino acid sequence will be cut into units with either triple- or tetra-amino acids (Figure. 2A). Next, a script for scoring system based on the structure of group I intron, and the rule of degeneracy of genetic codes was developed. Simply, the scoring method is according to the rule developed by Rausch et al, each nucleotide will be judged and scored as 0, 1, 2 or 3 points, depends on whether a nucleotide is identical as the one in specific position of T4td E1-E2, and whether it can pair with the specific nucleotide of internal guide (IG) sequence of T4td intron. Finally, all possible amino acid units will be judged, and all the potential segments will be scored, from 0 to 16 points. The segments with higher scores are predicted to conduct better circularization.

**Figure 2:**
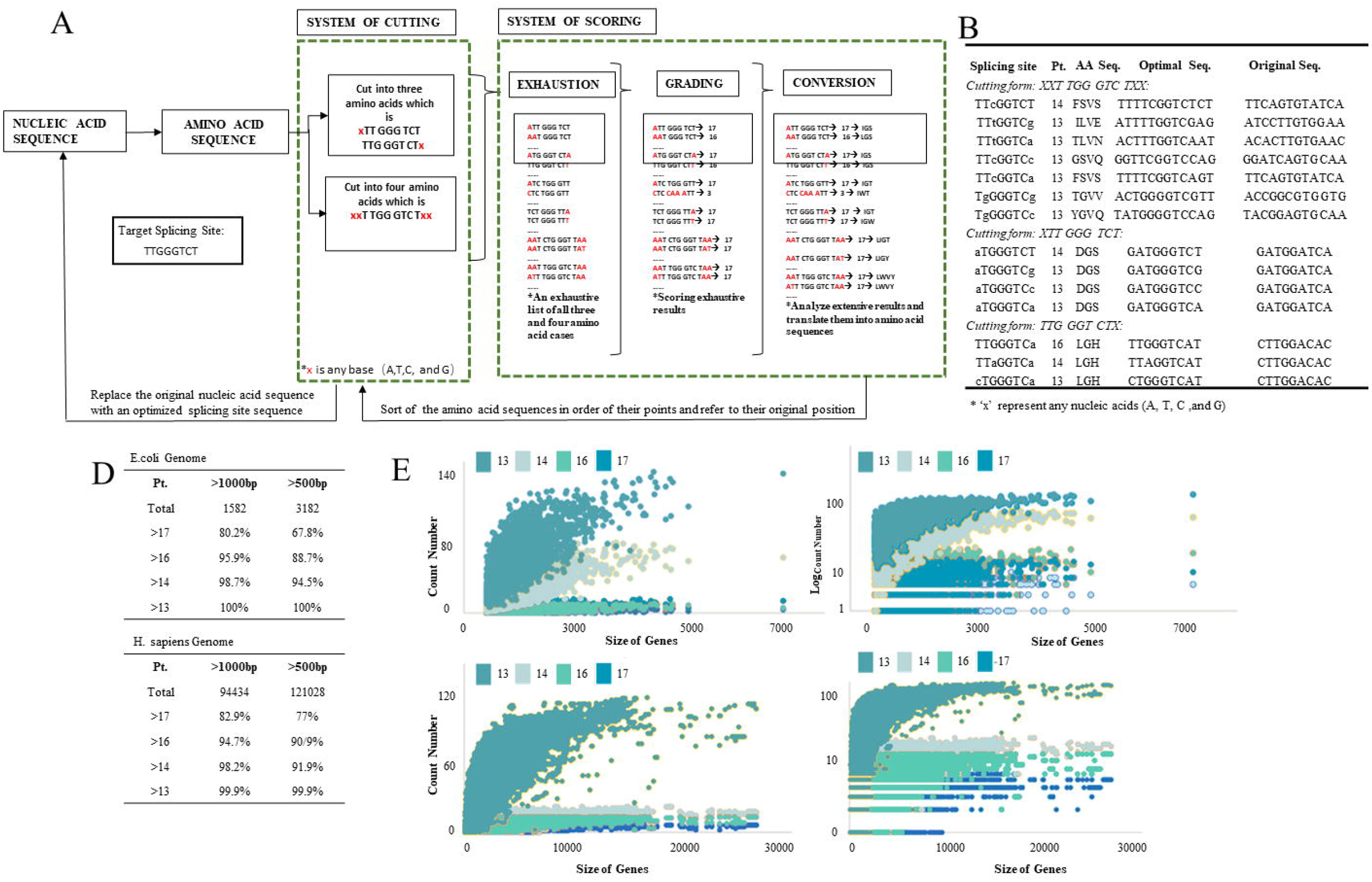
A novel scoring method is used to determine the splicing site automatically in ORFs. A) A diagram illustrating the pipeline for the automatic splicing site concealing system. The system is exampled with the *Thymidylate* synthase intron. B) The optimal splicing site in eGFP is automatically identified and listed. Pt: point, AA seq: amino acid sequence, Optimal seq: optimal sequence, Original Seq: original sequence. C) A statistic detailing the number of optimal sites in E. coli and H. sapiens over a period of 13 points. Pt: points. D) Scatter plots of the statistic of the optimal sites which above 13 points in E. coli and H. sapiens.

With this automatic script, the sequence of eGFP is revisited. Three additional splicing sequences were found with only single different nucleotide compared to the E1-E2 sequence of the original T4td PIE (Figure. 2B). Finally, the selected optimal segments and its corresponding amino acid units were identified, consequently, DNA construct sequences for linear RNA precursor, with elements including the flanked introns, segmented ORF and IRES, can be generated, then circRNAs were produced after transcription and circularization. To test whether this automatic script can be generally used, all E. coli and H. Sapiens genes were evaluated with this system. As shown in Figure. 2C and 2D, 80.2% of the E. coli genes with the size >1000bp contain at least one segment with the same sequence as T4td PIE E1-E2 (‘TTGGGTCT’). Even for genes with the size >500bp, 75% of total genes contain ‘TTGGGTCT’. Additionally, all 3182 genes that are larger than 500bp were found to contain at least one sequence with the score above 13 points, which means only single nucleic acid differs from the original E1-E2 sequence. 82.9% of the H. Sapiens gene with the size >1000bp contain at least one segment with the same sequence as T4td PIE E1-E2 (‘TTGGGTCT’), for genes with the size >500bp, 77% of total genes contain ‘TTGGGTCT’. 99.9% of the genes with the size >500bp or >1000bp contain at least one segment scoring as 13 points. In summary, these statistical results suggest that our circularization strategy with ORF E1-E2 segments probably is a general method for the construction of translational circRNAs.

### A variety of RNAs are effectively circularized with Clean-PIE(ORF)

The engineering *Anabaena* group I intron PIE invented by Wesselhoeft RA. *et al*. has demonstrated that different sizes of RNA precursors can be circularized^8^. Circularization of Gaussia luciferase (total length: 1289 nts), firefly luciferase (total length:2384nt), eGFP (total length: 1451 nts), human erythropoietin (total length: 1313 nts), and Cas9 (total length: 4934 nts) with different IRES have shown to be possible. Next, as a means of assessing the ability of this automatic splicing site concealing system to circularize different RNAs, we attempted to construct RNAs with different sizes for encoding different proteins by our method. As shown in Figure. 3A, the splicing sites with highest score are found in the sequences of murine IL12 and spCas9. However, splicing sites with only moderate scores, like 13 points (pts) and 14pts are found in Fluc and FLAG-con1-SPOP_167-274_^20^. Except for FLAG-con1-SPOP_167-274_, all other RNAs circularized efficiently, as determined by agarose gel analysis (Figure. 3B). The splicing site that is too closed to IRES may have contributed to unsuccessful circularization of FLAG-con1-SPOP_167-274_. This is consistent with the conclusion from Wesselhoeft RA. *et al*. that highly structured intron and IRES are not able to fold and function independently^8^.

**Figure 3:**
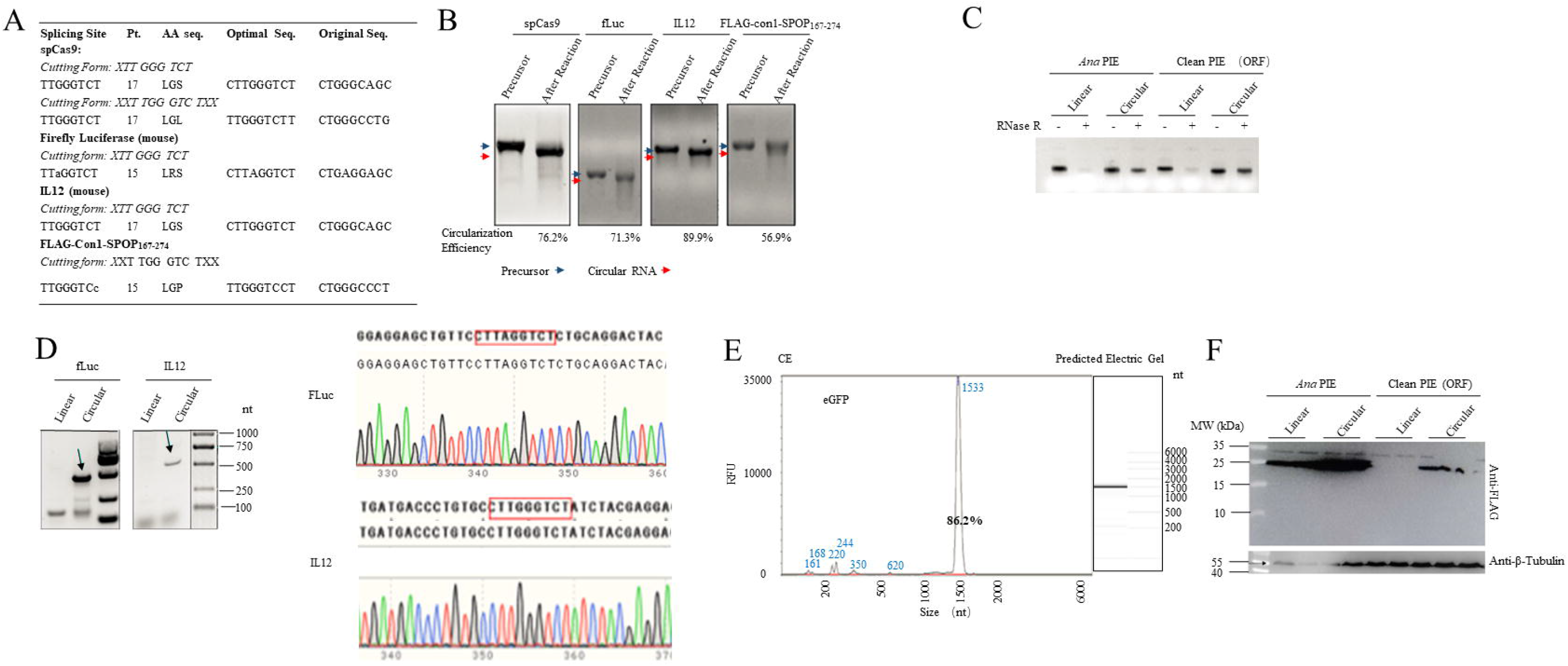
Clean-PIE (ORF) conducted effectively circularization of various RNAs. A) A table provides information regarding optimal splicing sites and splicing site sequences used for RNA circularization. Pt: point, AA seq: amino acid sequence, Optimal seq: optimal sequence, Original Seq: original sequence. B) The agarose gel electrophoresis demonstrates that circularization of different RNAs by Clean-PIE (ORF) is effective. A grayscale morphology analysis is used to measure the efficiency of circularization. Main bands of the circular RNA proportion in C) Linear RNA precursor and circRNA were treated with RNase R D) Sequencing of RT-PCR products of circRNA indicates a precise and uniform splice junction. E) A representative capillary electrophoresis analysis of circular eGFP is shown, along with the percentage of circular RNA bands. An example of electric gel is shown in the right-hand side of the figure. F) Western blotting analysis indicated that translation products cannot be generated using the linear precursors in Clean-PIE (ORF) strategy.

Additionally, as illustrated by Figure. 3C, RNase R digestion was performed to verify the circularization, since circRNAs, but not their linear RNA counterpart, are resistant to RNase R. This was further verified by the results of reverse-transcription polymerase chain reaction (RT-PCR) and Sanger sequencing, which indicated that a successful ligation of E1-E2 segment that concealed in ORF, suggesting that our automatic splicing site concealing system affords with high circularization effectivity (Figure. 3D). Our further studies revealed that the small fragments (introns) can be removed by HPLC-SEC, and thus resulted in around 90% purity of the product, as determined by capillary electrophoresis, suggesting a relative high circularization effectivity of our method (Figure. 3E). Taken together, all these results suggest that our automated splice site concealing system achieves efficient circRNA formation.

Cap-independent translation mediated by IRES is the most common choice in currently circular mRNA technology. However, in previous studies by using the ORNA circRNA constructing format, the linear precursor also exhibits varying levels of protein expression (data not shown), despite that the protein expressed by linear precursor in ORNA format is presumed to be the same as that expressed by circRNAs. However, the segmented and flanked of ORFs in our circularization format results in truncated ORFs in RNA precursors, raising a concern that RNA precursors may express truncated proteins. Thus, circRNAs and its linear precursor encoding N-terminal FLAG-tagged con1-SPOP_167-274_ were used to test this hypothesis, since the N-terminus of this protein is located following IRES, and both the full length or truncated form of this protein can be monitored by anti-Flag antibody in Western blotting assay. As shown in Figure. 3F, the Flag-tagged protein was found to express in circRNA, however, was not observed in the linear precursor, suggesting that RNA precursor did not express any protein. In comparison with this, protein expressed in the linear precursor of *Anabaena* group I intron PIE (ORNA format) was observed, with the same protein size as the one expressed by circRNA (Figure. 3F).

### Protein production is comparable to other circRNAs produced from different *in vitro* strategies

It is worth taking note that circRNAs that generated by our format led to relative lower protein expression, compared to the one from ORNA format, as revealed from FLAG-con1-SPOP_167-274_, (Figure. 3B), eGFP and murine IL-12 circRNAs. Comparing the final sequence of circRNA with engineered *Anabaena* group I intron PIE and our format indicated that the additional spacer sequences located between *Anabaena* E1-E2 and the coding sequence may be critical for the promotion of protein translation This suggests that the spacer sequences and Anabaena E1-E2 sequences contain some elements that can increase the translation activity of circRNAs. The spacer sequence includes a piece of polyAC sequences and part of the internal homology arm sequences. We suspected the polyAC sequences resemble the sequences of polyA tail of mRNA, which can be bound by factors such as polyA binding protein (PABP), thus to trigger the binding of the eukaryotic initiation factor 4 complex (eIF4G) to facilitate ribosomal initiation^21^. Different length of polyAC from 1 x (19 nts) to 10 x (190 nts) were investigated and tested for the expression of eGFP and murine IL12 (Figure. 4A, 4B). We found that polyAC increased the protein productivity of circRNAs in a length dependent manner, and 6x polyAC (around 114 nts) was found to be the optimal sequence to facilitate protein production, and even led to higher protein expression than the one from circRNA of ORNA format. However, 10x polyAC failed to induce higher protein expression (Figure 4B), this may be ascribed to a damage of translatability of circRNA by a too long polyAC sequences. Wesselhoeft RA *et al*. found that the spacer-spacer complementarity and the pairing of homologous arms formed a sheltered splicing bubble potentiates the catalytic intron splicing. As a consequence of concealing the splice site in the coding sequence, it is hard to form such a splicing bubble in our RNA construct. However, probably the paring of homology arm still contributes to RNA circularization in our format. We tested the circularization effectivity of RNA constructs with short or long homology arms. As shown in Figure. 4C, all these designs are circularized effectively, suggesting that the homology arms play a limited role in the effectivity of RNA circularization in our format.

**Figure 4:**
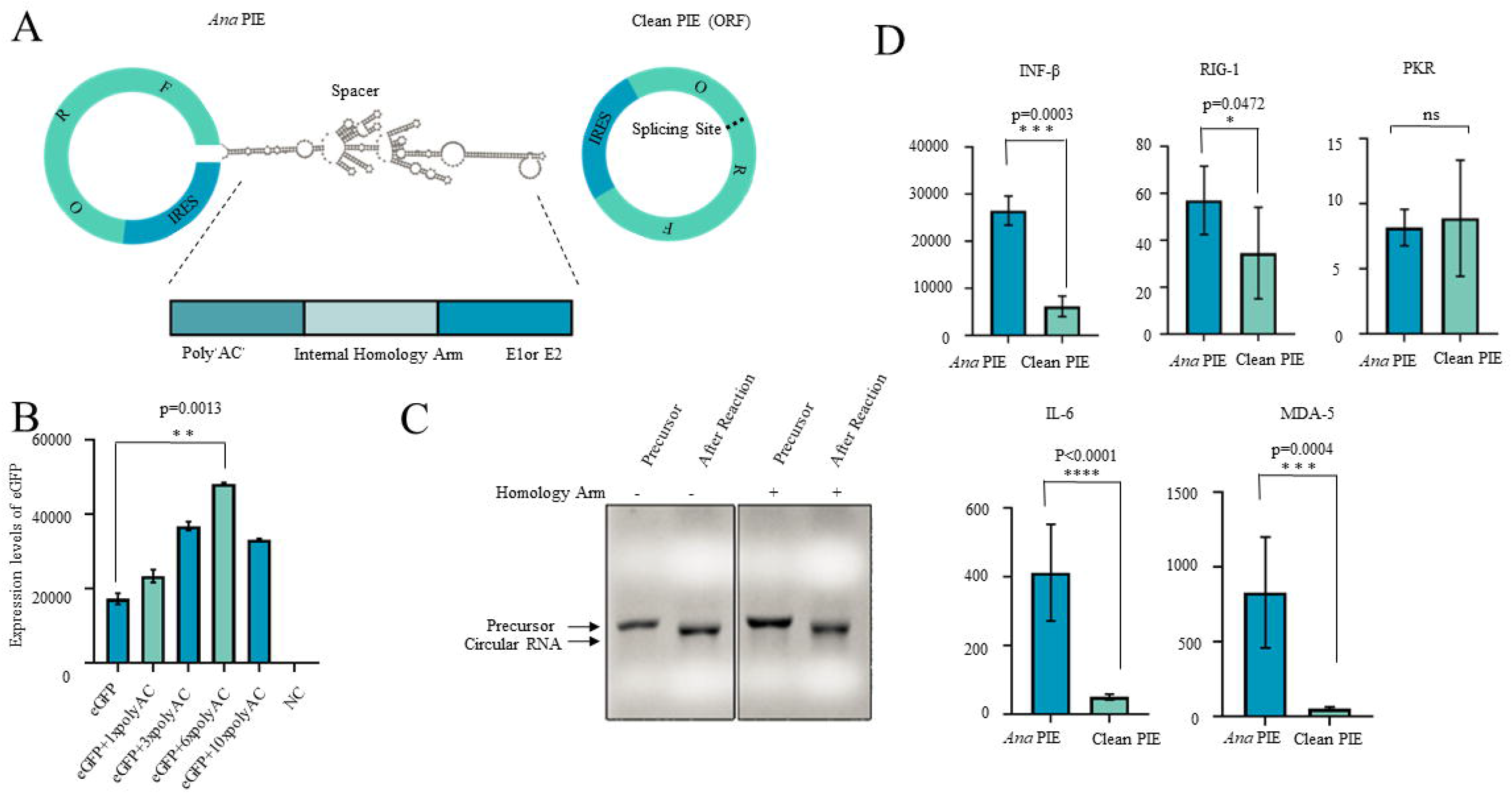
RNA is effectively circularized by Clean-PIE (IRES) A) A schematic diagram illustrating the difference between the final product of Ana PIE and Clean-PIE. A description of the spacer composition is given in the schematic diagram. B) The polyAC tracts attached to the eGFP ORF exhibit the highest degree of expression. C) Agarose gel analysis demonstrated that circularization efficiency of RNA is not affected by the absence of the homology arm. D) Circular RNA generated *via* Clean-PIE is associated with minimal immunogenicity in comparison to that produced *via* Ana PIE. Expression levels of IFN-β, RIG-1, PKR, IL-6 and MDA-5 are quantified by real-time qPCR.

### Circular RNA produced via Clean-PIE (ORF) shows thoroughly minimized immunogenicity

Previous studies have reported that circular RNAs made by different *in vitro* circularization strategies can induce distinct innate immune responses^12^. As concealing the splicing site into the ORF, circRNAs that are much more precise are produced, and extraneous RNA sequences were not involved in. In the current study, we examined the immunogenicity of thoroughly purified (agarose gel separation) circRNAs that generated by Clean-PIE (ORF) and *Anabaena* PIE (ORNA format). The transcription levels of serval inflammatory cytokines and innate immune regulators were measured after transfection of circRNAs in A549 cells. Firstly, CircRNA POLR2A is constructed by both *Anabaena* PIE (circPOLR2A_*Ana* PIE) and our Clean-PIE(ORF) (circPOLR2A_Clean-PIE)^12^. The same amounts of circPOLR2A which were produced by different strategies and were purified by agarose-gel, were transfected into A549 cells, and the expression of IFN-β, RIG-1, MDA5, IL-6 and PKR were measured at 6 and 24 h post transfection. As shown in Figure. 4D, compared to the group of circPOLR2A_*Ana* PIE, much lower immunogenicity was observed in the group of circPOLR2A_Clean-PIE, characterized by the thorough low expression of IFN-β, IL-6 and MDA-5 in Clean-PIE circRNA group. Conversely, differences were not observed in PKR, this may be explained by the fact that PKR is distinct from TLRs and RLRS in the mechanism of actions, PKR appears to directly target RNA either through inhibition of translation or global destruction of RNA, respectively^22,23^. Taken together, these results suggest that the avoid of introducing extraneous RNA sequences from Anabaena or T4td into circRNAs brings to thoroughly minimized immunogenicity of circRNA.

### T4td splicing site concealing in translation initiation sequence (IRES) resulted in high circularization effectivity and potent protein expressions

According to the report from Weesselhoeft et al., the translation initiation sequence, or exactly IRES of Coxsackievirus B3 (CVB3) was identified as the most potent IRES for the direction of circRNA mediated protein expression. In our previous reports, we identified the IRES of Echovirus 29 (E29) and E33 as stronger element to induce protein expression^24^. In the current study, we broadened the screening of IRES in varying strains of virus and attempted to identify better IRES for Clean-PIE circRNA. More than 600 IRES elements were selected by an algorithm assisted virtual screening method (Figure.5A), and were inserted to the ORNA circRNA framework that encoding *Fluc* for the evaluations of translational activity. The circRNA library with various IRES elements was screen by firefly luciferase activity, and more than 20 IRES candidates were identified to mediated significant higher translational activity than E29, as illustrated by Figure. 5A. Furthermore, we tried to analyze the potential segments in these IRES elements, and found several of them possesses ideal segment scores, as judged by our automatically scoring system. Thus, IRES309 was selected as the candidate for E1-E2 concealing, and thus were segmented and flanked besides the two introns in the novel circRNA precursor constructs, and the IRES309 E1-E2 was ligated by autocatalytically ribozymatic reaction, as shown by Figure. 5B. Similar with the Clean-PIE (ORF) method, this strategy won’t introduce extraneous RNA sequences, thus are termed “Clean-PIE (IRES)”. Next, circRNAs encoding murine IL-12, eGFP, FLuc, Flag-tagged con1-SPOP_167-274_ were designed, and circularization reactions were performed. RNAs that after circularization reactions were analyzed by denaturing agarose gel electrophoresis, as revealed by Figure. 5C, all of the four RNA precursors were circularized, with the effectivity of 88.2%, 89.1%, 82.5% and 81.7%, respectively. For further validation of circularization, precursors and circRNAs were both treated by RNase R, which selectively digest linear RNA. Results of Figure. 5D ascertained that the four circRNAs were resistant to RNase R, compared with the precursor counterparts. Moreover, the circularization reactions were determined again by the digestion of RNase H, which digest the DNA-RNA hybrid strands. Thus, after the binding of DNA probes to the E1-E2 junction of IRES309, RNase H digested the circRNAs to generated one strand of open-ring RNAs, of which sizes can be determined by the mobility in agarose gel. As revealed by Figure.5E, RNase H digestion of precursors led to two strands of RNAs, however, yield a single strand for circRNAs. Finally, the circularization was validated by RT-PCR and Sanger sequencing, as illustrated by Figure.5F, the intact sequences of IRES segments were found in the circRNA groups, but not in precursor, suggesting successful ligations of the segmented IRES to form circRNAs. Next, comparisons of circRNAs mediated eGFP and Flag-tagged con1-SPOP_167-274_ protein expressions among the three circularization construction formats, including Clean-PIE (ORF), Clean-PIE (IRES) and Ana PIE (ORNA format), were performed. As indicated by Figure.5G, slight eGFP protein expressions were found in the group of cells that transfected with RNA precursors of Clean-PIE (IRES) and Ana PIE, but not in Clean-PIE (ORF), potent eGFP protein level was found in the group of all the three circRNAs; moderate con1-SPOP_167-274_ protein expressions were found in Ana PIE precursor and Clean-PIE (ORF) circRNAs, however, potent expression in Ana PIE and Clean-PIE (IRES). These results suggested that circRNAs that generated by Clean-PIE (IRES) elicited protein expression that almost identical to the circRNAs of Ana PIE.

**Figure 5:**
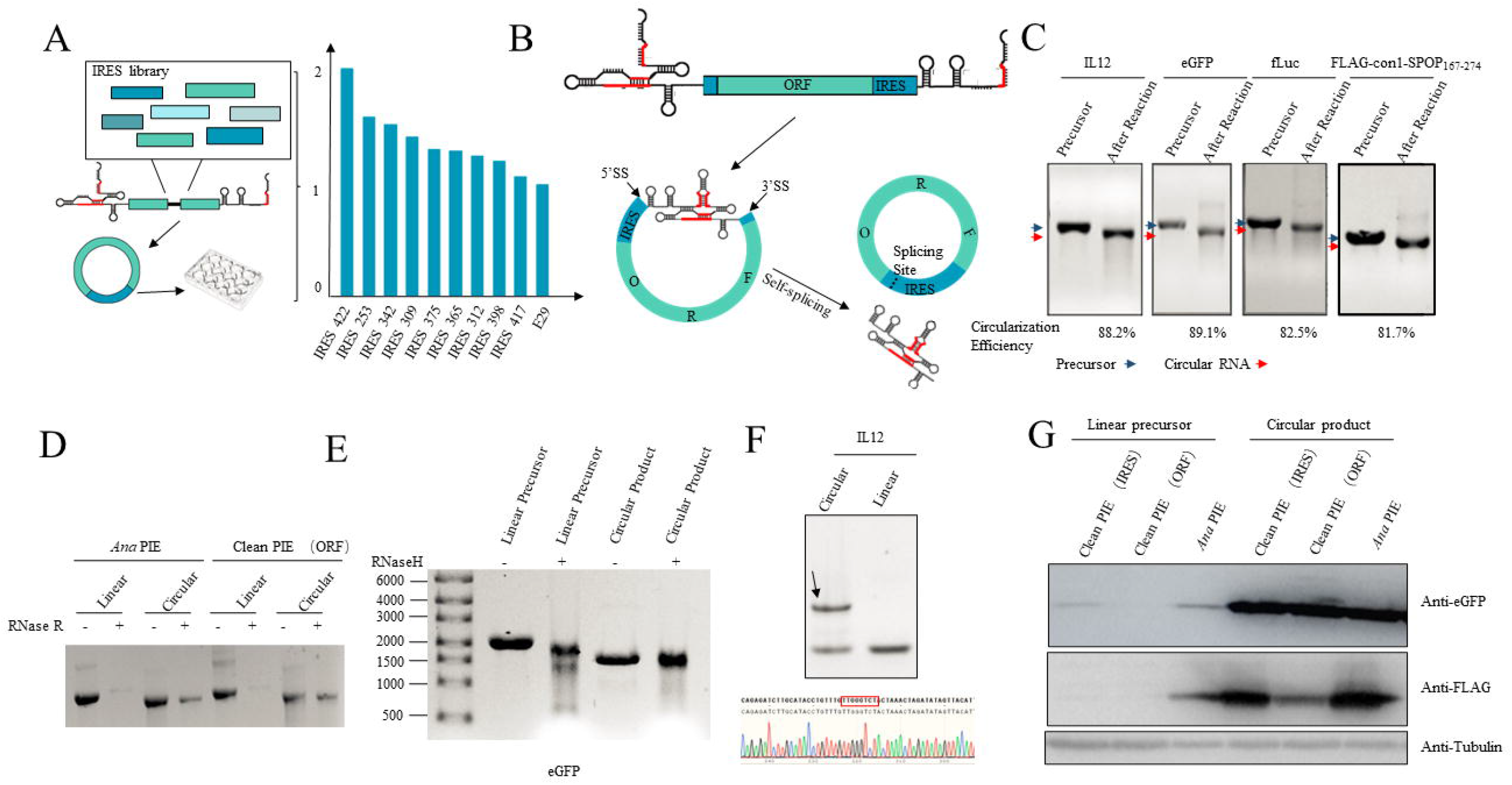
Immunogenicity of circular RNA produced by splicing site concealment is minimized. A) A schematic overview of the procedure to identify effective IRES. B) A diagram showing the design of a permuted intron-exon construct that conceals the IRES splicing site. C) The agarose gel demonstrates that PIE circularization at the IRES of different RNA is effective. A grayscale morphology analysis is used to measure the efficiency of circularization. Main bands of the circular RNA proportion. D) Linear precursor RNA and circular RNA were treated with RNase R to prove the RNA is circularized efficiency. E) Linear RNA precursor and circular RNA were treated with RNase H to prove the RNA is circularized efficiency. F) Sequencing of RT-PCR products of circRNA indicates a precise and uniform splice junction at the IRES of the circRNA. G) Expression products of both precursors and circRNAs from *Ana* PIE, Clean-PIE (ORF), and Clean-PIE (IRES) are analyzed, which is conducted using western blot.

### Improved circRNA purification leads to higher protein expression and scaling-up in Clean-PIE (IRES) format is accessible

As described by Wesselhoeft RA. *et al*, PIE mediated circularization often leads to side products, such as the small RNA fragments (introns) generated by splicing, the remnant of linear RNA precursors which is from the failure of circularization, and the nicked RNAs, which is the linear isoform of circRNAs (PMID). Thus, the development of RNA purification methods may be critical to obtain high-purity circRNA. In our previous studies, we found that the small intron fragments can be removed by HPLC-SEC method^25^. Here, we provided evidence to show that the eGFP circRNA that were generated by Clean-PIE (IRES) or Ana PIE format, can be further purified by a new purification method to remove the small intron fragments, the remnant of RNA precursors and the nicked RNAs, and the purity was determined as around 90% by Urea-PAGE (Figure. 6A). Next, the high-purity circRNAs were found to mediate much more potent eGFP protein expressions in A549 than the ones purified by HPLC-SEC, in which only free-style intron fragments were removed (Figure.6B). Next, we explored the protein expression duration mediated by Fluc circRNAs that generated by Clean-PIE or Ana PIE format in muscle, by direct injection of naked, HPLC-SEC purified (free-style introns removed) circRNAs. As revealed by Figure.6C, the expression of FLuc in muscle by circRNAs from either Clean-PIE (IRES) or Ana PIE, started with 1× 10^6^ of fluorescence flux, decreased gradually to around 1× 10^4^ of fluorescence flux at the day 20, suggesting an extreme long-duration of protein expression by circRNAs in vivo. Next, the comparison of Fluc protein expression in muscle by among circRNAs of Clean-PIE (IRES) with HPLC-SEC purification and the new purification method. As shown in Figure.6D, the expression by circRNAs that purified by our new method exhibited significant stronger Fluc expression than the one purified by HPLC-SEC, and reached to a top value at 48 h, and decreased mildly at 72 h. The studies of protein expression duration by high-purity circRNAs are still ongoing, we prospect a much better performance than circRNAs that purified by HPLC-SEC.

**Figure 6:**
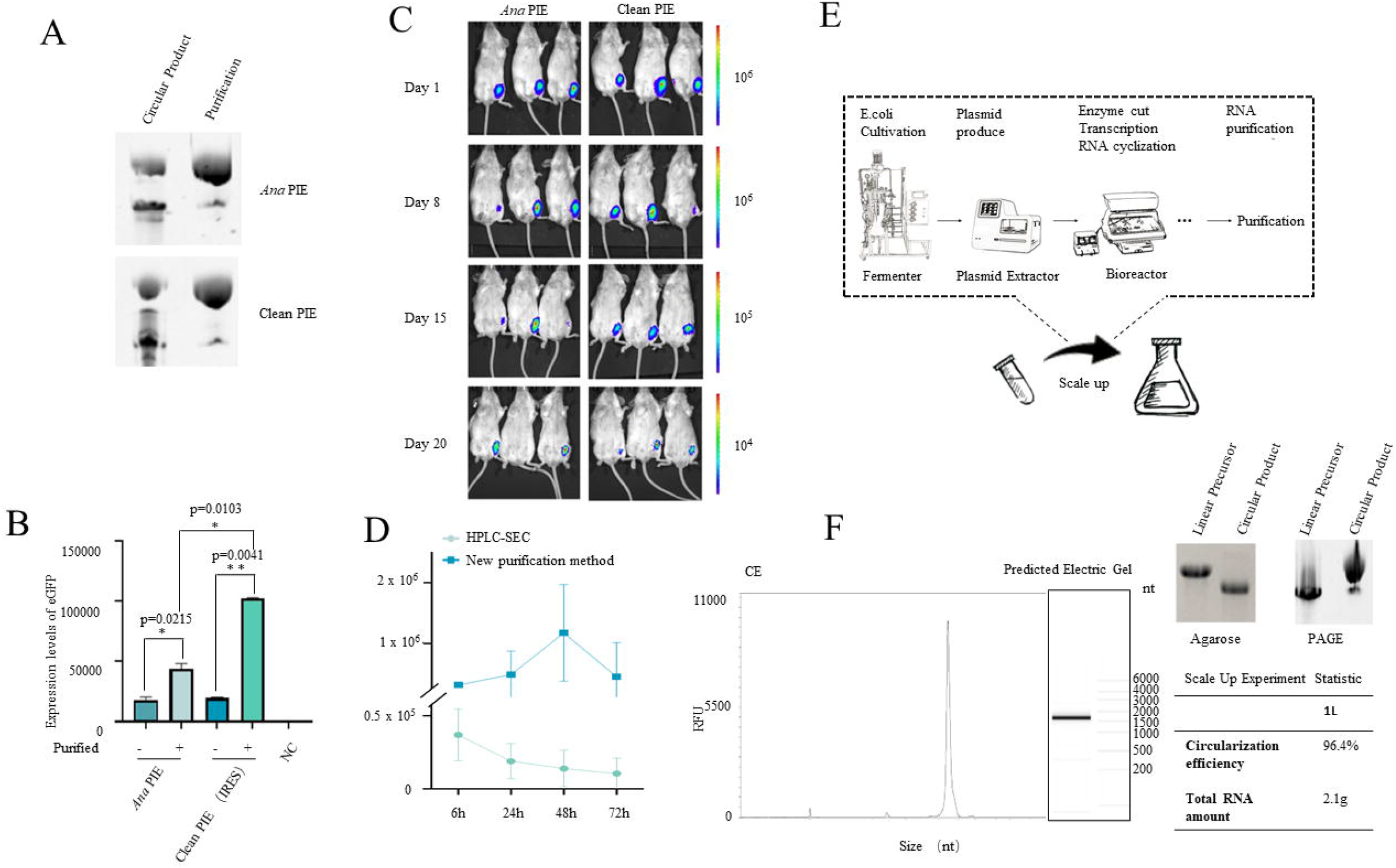
New purification method for circRNA improved protein expression and scale-up of circRNA manufacturing by Clean-PIE (IRES) A) High-purity circRNA is obtained by the new purification method. The purified circRNA was analyzed by Urea-PAGE. B) CircRNA directed eGFP expression levels in A549 have dramatically increased after purification by the new method, compared to the HPLC-SEC method. C) A series of IVIS images taken of BALB/c mice administered with 10 μg Fluc circRNA by ORNA format and Clean-PIE (IRES) format intramuscularly (i.m.) for long-term monitoring. D) CircRNA directed Fluc expression levels in the muscle of mice have dramatically increased after purification by the new method, compared to the HPLC-SEC method. E) A schematic diagram illustrating the scale up process of the Clean-PIE (IRES). Circularization effectivity in scale-up manufacturing was determined by capillary electrophoresis and agarose gel analysis. The table shows circularization effectivity and productivity of the scale-up manufacturing. Urea-PAGE was also performed to detect the purity of the circRNA that after purification.

Next, we investigated the scaling-up of Clean-PIE (IRES) circRNA manufacturing up to 1 L of transcription volume (Figure. 6E). Plasmids were prepared by fermenter, and linearized by restriction enzyme mediated digestion in bioreactor, and followed by 1 L scale in vitro transcription and circularization reaction in bioreactor (Figure. 6E). The transcription in 1 L volume of resulted in about 4 g of linear RNA precursor. The circularization effectivity was as high as around 96.4%, as determined by denaturing agarose gel electrophoresis (Figure. 6F) and capillary electrophoresis (Figure.6F). Next, the circRNAs were further purified by our new purification method to reach around 90% purity, as determined by Urea-PAGE (Figure. 6F). Finally, the total yield of circRNA after circularization was determined to be 2.1 g. These results suggest that the scale-up of Clean-PIE (IRES) circRNA is accessible and ready for large-scale manufacturing.

## Discussion

Although it has been demonstrated that circRNAs can be translated in a cap-independent manner with long half-life, and in vitro generated circRNAs for large scale and stable protein expression in eukaryotic cells have been realized recently^8^. The advanced property of circRNA, such as stable and directing long-duration of protein expression, makes it to be a preferred alternative to linear RNA^20,24–27^. Several circularization strategies have been proposed^28–31^. The most common methods for *in vitro* circularization of RNA are mainly chemical methods and enzymatic catalysis methods. Among them, the formation of natural phosphodiester bonds in the chemical circularization transformed the linear RNA precursor into a ring. However, chemical circularization has low ligation efficiency and is only suitable for ligating to generate small circRNAs, and chemical groups also have potential safety hazards. Therefore, chemical method for circRNA formation is considered to be inappropriate for widespread applications. Circularization by enzyme catalysis is mainly through T4 DNA ligase and T4 RNA ligases, as well as the recently identified RtcB enzyme^32^. However, the low ligation efficiency by these enzymatic methods may limit their potential applications. Additionally, the 3’heterogenicity of RNA precursors may also hinder the generation of ‘unalloyed’ and ‘precise’ circRNAs. The PIE method utilizes a permuted intron-exon system to conduct ribozyme mediated autocatalytic splicing reaction to generate circRNAs. This method has been proven to have high circularization effectivity and easy for scaling-up in industry, however, also processes disadvantages: i) the involvement of exon sequences from Ana or T4td may present immunogenicity, which may be a concern for the future applications of circRNAs as therapeutics; ii) generating circRNAs by PIE is easy to bring side-products, like the free-style fragments of introns that produced by splicing reaction, the nicked RNAs which are the open-ring isoform of circRNAs, thus it is difficult to obtain high-purity circRNAs. The aim of this study is to overcome the two disadvantages in PIE circularization method.

In the current study, we generate a new strategy to concealing splicing site into the ORF or translation initiation sequences (IRES). In our circRNA construction method, the segment sequences in ORF were identified in a given RNA/protein sequences according to the basic principle developed by Rausch. Et.al, and according to the degeneracy of genetic codes, the segment sequence can be optimized to an ideal one by replacing nucleotides, without a change for the corresponding amino acids. This method was found to conduct successful circularization with high effectivity. Next, a bioinformatic tool was developed to identify and optimize the segment sequences automatically, and this method was utilized to conduct circularization of a variety of circRNAs that encoding different proteins with high effectivity, demonstrating that it is a universal method for circRNA construction. Moreover, no protein expression was found from the linear RNA precursor, suggesting that the remnant of linear RNA precursor is not leading to the expression of truncation protein. Additionally, the incorporation of 6x polyAC sequences into circRNA increased its translation ability, which was defined to be equal to or even higher than the circRNA that constructed by ORNA format. Importantly, with this new circRNA construction method, we demonstrate that the avoid of incorporating extraneous sequences into the circRNA results in thoroughly minimized immunogenicity, as revealed by the detection of various innate immunity biomarkers, therefore, this method is termed Clean-PIE (ORF). This was also observed by Liu CX *et al*. with different circularization strategies^11^. CircRNAs that generated by *Anabaena* Group I intron PIE (Weesselhoeft *et al*.) contain extraneous sequences from *Anabaena* exons, which possesses dsRNA structures contributing to their innate immune responses. The elimination of immunogenicity of circRNAs by our method is of great importance, because: i) the immunogenicity of RNA leads to cellular responses to produce cytokines, like IFN-β, which results in a suppression of protein translation; ii) one of the most important properties of circRNA is its long half-life and duration of protein expression, the suppression of protein translation by RNA induced immune responses may impair the expression duration; iii) the damage of long protein expression duration by RNA immunogenicity may eliminate the advantages of circRNA in the potential applications in protein replacement therapies, circRNA directed CAR or TCR expression in ex vivo or in vivo CAR-T and TCR-T therapies, as well as circRNA directed gene modified stem cell therapies, etc.

For the development of Clean-PIE (IRES), we screened for over 600 IRES sequences and identified more than 20 IRES can mediate stronger protein expression than E29, which was confirmed to be stronger than CVB3 IRES in our previous study^25^. After searching for segment sequences of these IRES, we identified IRES309 as the one possessing ideal splicing site. The design of IRES309-segmented Clean-PIE (IRES) format results in highly effective RNA circularization (around 90% effectivity) for a variety of circRNAs that encode proteins with different sizes. The circularization was confirmed by a few methods, including agarose gel electrophoresis, Rnase R and Rnase H digestions. Importantly, circRNAs that constructed by Clean-PIE (IRES) have identical RNA sequences and structures as the ones constructed by Clean-PIE (ORF), therefore, it is rational to consider that they have the same thoroughly minimized immunogenicity. Additionally, Clean-PIE (IRES) method presented almost the same universality as Ana PIE, it means that this circularization format is theoretically compatible for any nucleic acid sequence that encodes any protein.

The circularization reaction by PIE methods leads to side products, including the remnant of linear RNA precursors, the free-style intron fragments after splicing, and the open-ring isoforms which were termed nicked RNAs. As linear unmodified RNAs, these side products are considered as strong immunogens, thus it raises a high requirement for thoroughly purification of circRNA products. In our previous studies, we found that it is easy to remove the free-style intron fragments by HPLC-SEC, due to their low molecular weights^25^. The removal of linear precursors and nicked RNAs is challenging, especially for large-scale industrial manufacturing. In the current study, we developed a new RNA purification method, with which the purity of circRNA can reach around 90%, as determined by Urea-PAGE. The high-purity circRNAs that generated by Clean-PIE (IRES) or Ana PIE exhibited much stronger protein expression in vitro, demonstrating the great importance of the removal of impurities. The in vivo durations of protein expression by naked circRNAs that generated by Clean-PIE (IRES) or Ana PIE, and purified by HPLC-SEC to remove free-style introns were observed. Both of the two Fluc circRNAs directed luciferase expression in muscle for at least 20 days, which is an extreme long-duration. The duration mediated by circRNA of Clean-PIE (IRES) did not show advantage against the one of Ana PIE, probably due to the limited purity of these two circRNAs, without removal of linear precursors and nicked RNAs, thus the overall immunogenicity by impurities may overwhelm the differences of the intrinsic immunogenicity between the two kinds of circRNAs. The comparisons of in vivo Fluc expressions between the circRNAs that purified by HPLC-SEC and by the new purification method proved that a strong facilitation of protein expression by removing the impurities, again demonstrating the importance of the purity of circRNA for protein expression.

Finally, we proved that the large-scale manufacturing of circRNAs by Clean-PIE strategy is accessible, evidenced by the successful circularization of the RNA precursors from 1 L transcription reaction, leading to high circularization effectivity and high product yield. The final circRNA product can be purified to be around 90% purity by the new HPLC method as well, demonstrating that our Clean-PIE based circRNA manufacturing platform is adequate for large-scale industrial manufacturing, and provides high quality circRNAs with thoroughly minimized immunogenicity and complete removal of impurities.

Collectively, we developed a novel circRNA construction strategy termed Clean-PIE, with splicing sites concealing in ORF or IRES, to generate circRNAs effectively that possessing thoroughly minimized immunogenicity and mediates potent and durable protein expressions in vitro and in vivo. The new purification method to obtain high-purity circRNA further reduces the final immunogenicity and promotes protein expressions. Additionally, the large-scale manufacturing based on Clean-PIE platform has been approved, which will enable the industrial development of circRNAs as the potential next generation of therapeutics and vaccines, as well as gene modification tools for cell therapies.

## Materials and Methods

### Human cell lines

Human cell lines including A549, and HEK293T cells were purchased from the American Type Culture Collection (ATCC; http://www.atcc.org)

### Cell culture and transfection

HEK293T and A549 cultured in Dulbecco’s modified Eagle’s medium (DMEM, BI) supplemented with 10% fetal calf serum (BI) and penicillin/streptomycin antibiotics (100 U/ml penicillin, 100 μg/ml streptomycin; Gibco) and maintained at 37 °C, 5% CO_2_, and 90% relative humidity. 1×10^5^ cells per well were seeded in 24 well plates or 4 ×10^5^ cells per well were seeded in 6 well plates. 500ng circRNAs per well for 24-well plates (2 μg for 6-well plates) were transfected with lipofectamine MessengerMax (Invitrogen, LMRNA003) on the next day when the cell culture must have >90% viability and be 70% confluent. Cells were collected after 24 h post-transfection for the following detections.

### Gene cloning and vector construction

DNA fragments containing PIE elements, IRES, coding regions, and others were chemically synthesized and cloned into restriction digestion linearized pUC57 plasmid vector. DNA synthesis and gene cloning were customized and ordered from Suzhou Genwitz Co., Ltd. (Suzhou, China).

### Plasmid extraction, *in vitro* transcription, and RNA circularization

CircRNA precursors were synthesized in vitro from a linearized plasmid DNA template using a Purescribe™ T7 High Yield RNA Synthesis Kit (CureMed, Suzhou, China). Following in vitro transcription, reactions were incubated with 15 minutes of Dnase I (CureMed, Suzhou, China). GeneJET RNA Purification Kit (Thermo Fisher) was used to column purify linear mRNA after Dnase I treatment. GTP was added to a final concentration of 2 mM along with a buffer that included magnesium (50 mM Tris-HCl (pH 8.0), 10 mM MgCl_2_, 1 mM DTT; Thermo Fisher). After heating RNA to 55°C for 15 min, RNA was column purified. RNA was separated on agarose gels using Bio-Rad’s agarose gel separation system. The ssRNA ladder (Thermo Fisher) was used for standard analysis.

### circRNA purification

A 30 * 300 mm size exclusion column with a particle size of 5 *m and pore size of 1000 * was used for high-performance liquid chromatography (HPLC) on a SCG protein purification system (Secure Instruments, Suzhou, China). A flow rate of 15mL/min was used when running RNA in RNase-free Phosphate buffer (pH:6). By detecting UV absorbance at 260 nm, RNA was detected and collected. After concentrating the purified circRNA in an ultrafiltration tube, phosphate buffer was replaced with RNase-free water.

### Protein expression analysis

After transfection, GFP fluorescence was observed and imaged using an Olympus IX70 Microscope Cutaway Diagram. In the 24 h following transfection, the average fluorescence intensity of the cells was determined by flow cytometry. HEK293T-GFP cells and HEK293T control cells were digested by trypsin and suspended in DMEM supplemented with 10% FBS and 1% penicillin/streptomycin. The cells were then washed twice and resuspended in PBS (Thermo Fisher) as directed by the manufacturer. The fluorescent signals were detected for 10,000 events on a BD FACScan.

### RNA isolation, RT-PCR and qRT-PCR

A total of RNA was extracted from cell-cultured samples using Trizol (Life Technologies) in accordance with the manufacturer’s protocol. The extracted RNA was then treated with DNase I (CureMed, Suzhou, China). The cDNA was reverse transcribed and applied for PCR/qPCR analysis with PrimeScript™ reverse transcriptase (2680B, Takara, Japan). As an internal control for normalization, actin was examined. Three independent experiments were conducted to determine the expression of each gene.

### RNase R and RNase H analysis

The RNase R was digested following the manufacturer’s instructions (BioVision), 1μg of linear or circular RNA was digested with 1U of RNase R for 15-30 min at 37°C. The enzyme was inactive at 65°C for 10 min. For RNase H nicking analysis, 1:5 ratio of RNA: DNA duplexes were first heated at 65°C for 5 min. And then annealed at room temperature for 10 min. Reactions were incubated with RNase H (New England Biolabs) at 37°C for 10 min and stopped with the addition of 1μl of 0.5M EDTA.

### Urea-PAGE

For Urea-PAGE, RNA was run in 4% denaturing urea polyacrylamide gel electrophoresis employing 7M urea. Before running the gel, high voltage power supply (Bio-Rad) was applied with constant watts (25W) for 20-30 min. RNA sample were mixed with 2 x loading buffer contains 90% formamide, 0.5% EDTA, 0.1% xylene cyanol and 0.1% bromphenol blue.

### Identifying potential splicing sites in ORF and automatic segment sequence optimization

The python script for finding the splicing site automatically and codon optimizing is provided in Supplementary Data. The screening process is following the diagram illustrated in Figure. 2A. A ‘.txt’ format file will be generated with different amino acid splicing forms. The score, original sequence, and optimal sequence will be found in the file. Once the optimal splicing site was identified and the sequence encoding protein was treated as described above for circulation.

### Statistic detailing the number of optimal sites in E. coli and H. sapiens

The genome of E. Coli and H. Sapiens was download from NCBI (https://www.ncbi.nlm.nih.gov/). All genes above 500 bp or 1000 bp were analyzed with the score.py obtained from Rausch JW^19^ which was attached to the Supplement materials. The results were counted with a counting script and the points of the sequences were statistics and shown with a scatter plot.

### Statistical analyzes

For comparisons of data at a single time point, the two-tailed Student t-test was utilized to determine statistical significance. For data comparisons with multiple time points, two-way ANOVA was used to determine the statistical significance. Statistical significance for the comparison of survival curves was determined using the Mann-Whitney log-rank test. Graphs and statistical analyzes were generated using Prism version 8.0 (GraphPad). Statistical significance is denoted as ns, not significant, *P < 0.05, **P < 0.01, and ***P < 0.001.

